# Inflammation promotes aging-associated oncogenesis in the lung

**DOI:** 10.1101/2024.03.01.583044

**Authors:** Catherine Pham-Danis, Shi B Chia, Hannah A Scarborough, Etienne Danis, Travis Nemkov, Emily K Kleczko, Andre Navarro, Andrew Goodspeed, Elizabeth A. Bonney, Charles A. Dinarello, Carlo Marchetti, Raphael A. Nemenoff, Kirk Hansen, James DeGregori

**Affiliations:** Departments of Biochemistry and Molecular Genetics, University of Colorado Anschutz Medical Campus, Aurora, CO, United States; Biomedical Informatics,University of Colorado Anschutz Medical Campus, Aurora, CO, United States; Immunology and Microbiology, University of Colorado Anschutz Medical Campus, Aurora, CO, United States; Medicine, University of Colorado Anschutz Medical Campus, Aurora, CO, United States; University of Colorado Cancer Center, University of Colorado Anschutz Medical Campus, Aurora, CO, United States; Department of Obstetrics, Gynecology and Reproductive Sciences, Larner College of Medicine, University of Vermont, Burlington, VT, United States

**Keywords:** lung cancer, inflammation, microenvironment, aging

## Abstract

**Background:** Lung cancer is the leading cause of cancer death in the world. While cigarette smoking is the major preventable factor for cancers in general and lung cancer in particular, old age is also a major risk factor. Aging-related chronic, low-level inflammation, termed inflammaging, has been widely documented; however, it remains unclear how inflammaging contributes to increased lung cancer incidence.

**Aim:** To establish connections between aging-associated changes in the lungs and cancer risk.

**Methods:** We analyzed public databases of gene expression for normal and cancerous human lungs and used mouse models to understand which changes were dependent on inflammation, as well as to assess the impact on oncogenesis.

**Results:** Analyses of GTEx and TCGA databases comparing gene expression profiles from normal lungs, lung adenocarcinoma, lung squamous cell carcinoma of subjects across age groups revealed upregulated pathways such as inflammatory response, TNFA signaling via NFκB, and interferon-gamma response. Similar pathways were identified comparing the gene expression profiles of young and old mouse lungs. Transgenic expression of alpha 1 antitrypsin (AAT) partially reverses increases in markers of aging-associated inflammation and immune deregulation. Using an orthotopic model of lung cancer using cells derived from EML4-ALK fusion-induced adenomas, we demonstrated an increased tumor outgrowth in lungs of old mice while NLRP3 knockout in old mice decreased tumor volumes, suggesting that inflammation contributes to increased lung cancer development in aging organisms.

**Conclusions:** These studies reveal how expression of an anti-inflammatory mediator (AAT) can reduce some but not all aging-associated changes in mRNA and protein expression in the lungs. We further show that aging is associated with increased tumor outgrowth in the lungs, which may relate to an increased inflammatory microenvironment.

## INTRODUCTION

Modern society has seen a steady increase in life expectancy in the last two-hundred years^1^. Thus, it is not surprising that elderly people (>65 years old) are expected to comprise greater than 20% of the world’s human population by 2050^2^. This increase poses a forthcoming challenge for the public healthcare systems, particularly in the management and treatment of cancer patients. Lung cancer is the leading cause of cancer-related deaths in the world. Smoking has been known to be the major risk factor in lung cancer. Still, ∼25% of lung cancers occur in never smokers^3^. While the etiology of lung cancer in never smokers is complex, aging is one of the most prominent prognostic factors with the median age of lung cancer diagnosis at 70 years^4^.

Aging is a complex process due to stochastic, environmental, and genetic events that affect multiple cell types, which ultimately results to declining tissue function. There are a number of aging-associated alterations that occur in the lungs of mice and humans, including inflamm-aging^2,5^, declining adaptive immunity^6,7^, weakening of muscles, and declining lung function^8^. Inflammation has long been connected to the promotion of cancer^9^. In fact, chronic obstructive pulmonary disease (COPD), a chronic inflammatory lung disease, is an independent risk factor for lung cancer^10^, and smokers susceptible to COPD are five times more likely to develop lung cancer compared to those with normal lung function^11^. In fact, COPD is highly associated with lung cancer risk even in never-smokers^12^, although there are many complicating factors that prevent a clear causative link^13^. Interestingly, COPD usually manifests late in life and shares many similarities seen in the aging lung^10,14^.

It is widely accepted that tumorigenesis is a result of clonal evolution, where genetic variability brought about by mutations and epigenetic changes fuels selection for cell clones with progressively more malignant phenotypes. Much of the field has focused on mutations as the limiting factor in tumor development by conferring a fitness advantage in comparison to non-mutated cells. Based on this notion, cancer incidence is widely believed to be mostly limited by to the time-dependent accumulation of driver mutations. The relationship to aging is thought to mainly derive from the time required for sufficient driver mutations to occur within a single cellular clone^15–17^. Previous work from our lab has demonstrated that while mutations are crucial, they are not sufficient for cancer development, and that the impact of aging or other insults on selective pressures needs to be considered^18–20^. The Adaptive Oncogenesis model encompasses natural selection and microenvironmental changes to describe how selective pressure brought upon by changes in the tissue microenvironment may impinge on the selection of mutations that become adaptive in the evolved microenvironment.

Given the multitude of changes with advanced age, including inflammaging in the lungs, we hypothesize that aging-associated inflammation contributes to the selection of adaptive oncogene-initiated clones. To study how the tissue microenvironment may influence oncogenic-initiated clonal expansions, we use an orthotopic model of lung cancer whereby cells derived from EML4-ALK fusion-induced adenomas are implanted directly into lungs of mice. EML4-ALK fusions are prevalent in lung cancers of never-smokers, with a smaller fraction in those of smokers^21^. We have previously shown that reducing inflammation in an aged background via transgenic alpha1 anti-trypsin (AAT) can prevent oncogenic adaptation in B-cell progenitors, coinciding with reduced systemic inflammation^18^. Notably, patients with AAT deficiency, which results in degradation of the lung extracellular matrix, prematurely develop COPD and exhibit a 4-5 fold increased risk of lung cancer independent of smoking status^22^. Here, we show that both human and mouse lungs and lung adenocarcinomas/squamous cell adenomas exhibit hallmarks of inflammation in older ages, as well as alterations in extracellular matrix related gene expression. We further show that reducing age-associated inflammation can dampen the selection for EML4-ALK-driven clonal expansions in the lung.

## METHODS

### Mouse studies

Animals utilized were the following: human-AAT transgenic (AAT) mice^23^, *Nlrp3^-/-^* mice^24^, young and aged C57BL/6 mice. Young and aged C57BL/6 mice were provided by the National Institute of Aging (NIA). The *Nlrp3^-/-^* and AAT mice were maintained on the C57BL/6 genetic background. These strains of mice are designated “young” (2-5 months), or “old” (20-28 months) throughout this document. Both sexes of each of these strains were used for experiments. Oncogenic events were introduced by direct implantations of cancer cells derived from EML4-ALK fusion-induced lung adenomas^25^. The University of Colorado Institutional Animal Care and Use Committee (IACUC) reviewed and approved all animal experiments. All animal experiments were conducted in accordance with the NIH Guidelines of Care and Use of Laboratory Animals.

### Genotyping mice

The following PCR conditions were used to genotype each mouse strain:

AAT: 95°C for 2 minutes, 35 cycles of 95°C for 30 seconds, 50°C for 30 seconds, and 72°C for 30 seconds, followed by 72°C for 5 minutes. The AAT gene product produces a 450 bp fragment and ApoB which serve as an internal control produces a 324 bp fragment. A secondary confirmation using AAT nested primers was also performed, producing a 249 bp fragment.

Primer sequences for genotyping are listed in Table 1.

**Table 1.**
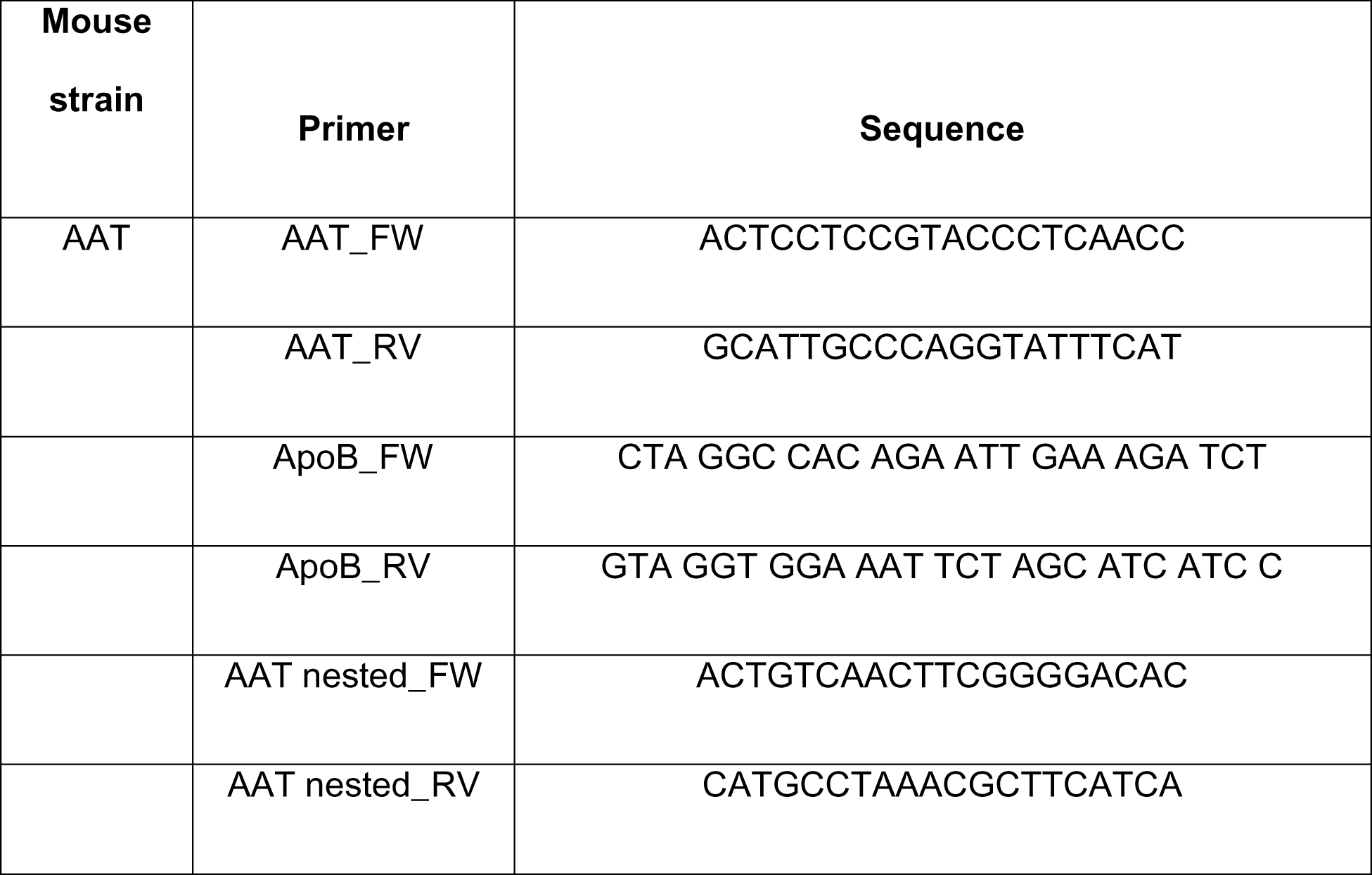
Primer sequences for genotyping.

### Preparation of slides and immunohistochemistry

Mouse lungs were evaluated through three serial 10μm sections obtained 50μm apart by hematoxylin and eosin (H&E) staining using standard method. Images were acquired using an Aperio Image Slide Scanner (Deer Park, IL, U.S.A.) or BZ-X810 inverted microscope (Keyence, Itasca, IL, U.S.A.). Images were analyzed using FIJI Image J software^26^.

### Lung tissue proteomics and analysis

Lung tissues from young, old, and old AAT mice were perfused with PBS and were pulverized. Lung tissues were then homogenized in freshly prepared high-salt buffer (50 mM Tris-HCl, 3 M NaCl, 25 mM EDTA, 0.25% w/v CHAPS, pH 7.5) containing 1x protease inhibitor (Halt Protease Inhibitor, Thermofisher Scientific) at a concentration of 10 mg/mL. Homogenization was performed using a bead beater (Bullet Blender Storm 24, Next Advance, 1 mm glass beads) for 3 min at 4 °C. Samples were then sequentially extracted using CHAPS buffer to obtain cellular and urea to obtain soluble fractions. The remaining pellet was then digested with either CNBr or NH_2_0H followed by tryptic digestion with QconCATs^27^. Samples were then analyzed by LC-MS/MS. MS/MS spectra were extracted from raw data files and converted into .mgf files using MS Convert (ProteoWizard, Ver. 3.0). Peptide spectral matching was performed with Mascot (Ver. 2.5) against the Uniprot mouse database (release 201701). Significant proteins were analyzed using Metascape (metascape.org).

### Lung epithelial cell gene analysis

Lung epithelial cell isolation, transcriptomic profiling, and analysis has been described elsewhere^28^. Briefly, lung tissues were homogenized using the MACS^TM^ (Miltenyi Biotec, Germany) tissue dissociation system, and CD45^+^ cells were removed using anti-CD45 (Miltenyi Biotec, Germany) beads. Affymetrix GeneChip system (Mouse Gene 2.0 ST Array) was used for transcriptomic profiling. Analysis of array data was carried out using Partek Genomics Suite® 6.6, Beta Analysis comparing groups with the number of probe sets passing an FDR of 0.05 or a binary filter (p<0.05 and 2× fold change).

### RNA seq lung preparation and analysis

RNA from lungs of young, old, and old-AAT mice was extracted from the lungs using Qiagen RNeasy mini kit according to the manufacturer’s protocol. cDNA library preparations were performed on the extracted RNA and were then run on the Illumina HiSeq at 50 cycles with 12 million read depth and FASTQ files generated. Low quality sequencing bases were trimmed before the alignment step. Trimmed sequencing reads were mapped against the mouse genome (mm10) using the Hisat2/cufflinks workflow. Cufflinks (version 2.2.1) were employed to assemble the transcripts using the RefSeq annotation as the guide and computed the transcripts’ RPKM values by using the merged assembly as the guide. On average, 40.4 million (28.8 – 59.2 million) reads were sequenced per sample and were aligned to the human genome. Gene Set Enrichment Analysis (GSEA) was used to perform pathways enriched in the RNA-seq as previously described. KEGG pathways were used as the gene set, and 1000 gene-set permutations were performed to determine p-value. Pathways with p<0.05 and logFC > |1| were considered significant from the analysis.

### GTEx and TCGA analyses

RNA-seq data were obtained from The Genotype-Tissue Expression (GTEx) Project GTEx portal on 08/04/2020. xCell^29^ analyses was done as described elsewhere. RSEM expected counts for LUSC and LUAD TCGA datasets (illuminahiseq_rnaseqv2-RSEM_genes) were downloaded from Firehose in January, 2022. Clinical data was downloaded as Firehose Legacy from cBioPortal^30^. The limma R package was used to calculate differential expression between the indicated groups^31^. Gene set enrichment analysis was performed using the fgsea R package on fold change using gene sets from the Molecular Signatures Database^32^.

### Orthotopic lung tumor model

EML4-ALK (EA) cell line was established by Kleczko et al, and the transplantation of EA cells into lungs of mice were described elsewhere^25^. Briefly, a 1mm incision was made to the left side of the anesthetized mice and 2×10^5^ EA cells resuspended in 40μl of Matrigel were injected directly into the left lobe of the lungs using 30-gauge needles, and the incision was closed with staples afterward.

### Tumor volume quantification

μCT imaging was performed by the Small-Animal IGRT Core at the University of Colorado Anschutz Medical Campus using the Precision X-Ray X-Rad 225Cx MicroIGRT and SmART Systems (Precision X-Ray, Madison, CT). ITK-SNAP software was used to quantify tumor volumes from μCT images^33^.

### Statistical analysis

For in vitro experiment, standard deviation is shown for all error and is based on biological replicates. Unless otherwise indicated, one-way ANOVA with Tukey post-test was used to compute p-values using GraphPad Prism. P-value designations for all figures: *<0.05, **<0.01, ***<0.001, ****<0.0001.

### Data Availability

The data supporting this study’s findings are available from the corresponding author upon reasonable request.

## RESULTS

### Advanced age leads to dramatic changes in the immune landscape of human lungs

While a dysregulation of the immune system has been reported to be a characteristic of old age^34^, we sought to evaluate the molecular and cellular changes that occur with advancing age in the lung. We analyzed the publicly available RNA-seq data from the Genotype-Tissue Expression (GTEx) project to determine gene expression changes in over 500 non-diseased lung tissue from individuals of various ages. We performed Gene Set Enrichment Analysis (GSEA) comparing the expression profiles of lung tissue from young (20-49 years old) to older ages (50-79 years old) to identify significant signatures (nominal p-value <0.05, p-value adjusted <0.2, and NES > 1.5 or < −1.5). We found that several pathways related to immune activation such as inflammatory response pathways, leukocyte intrinsic hippo pathway and T cell receptor and costimulatory signaling were enriched in the older compared to the young lungs (Fig. 1A). Additionally, we found that signaling pathways involved in inflammation including interferon gamma response and IL2/STAT5 signaling were also upregulated in the older samples relative to young (Fig. S1A). Extracellular matrix (ECM) remodeling pathways were also enriched in the older compared to young lungs suggesting alterations in the tissue architecture of old lungs (Fig. 1B). Interestingly, we identified Myc targets and oxidative phosphorylation gene sets enriched in young compared to older lungs (Fig S1B), which coincide with numerous studies showing a decline in Myc signaling^35^ and mitochondrial metabolism^36^ with advancing age.

**Figure 1.**
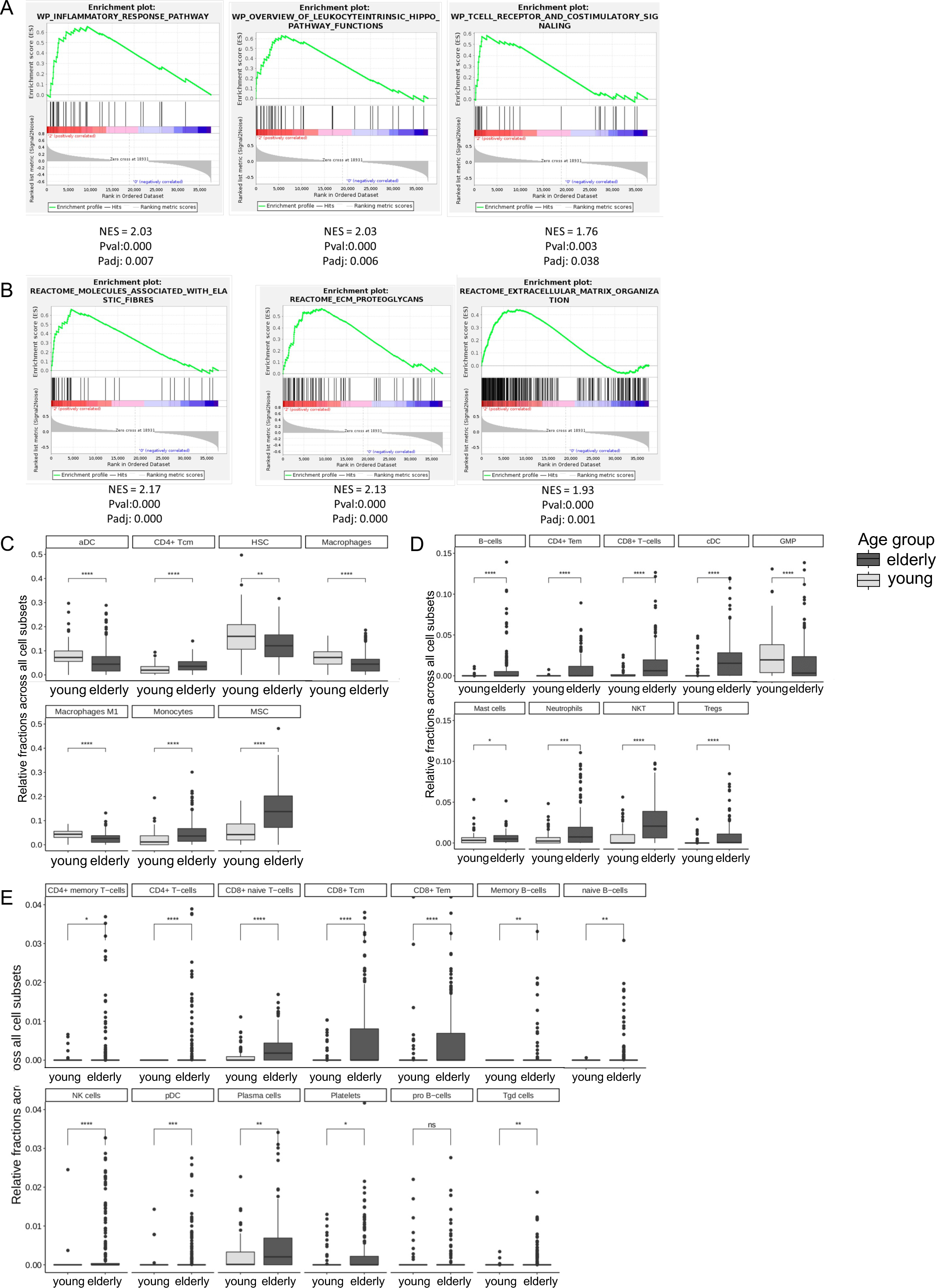
Advanced age leads to dramatic changes in the immune landscape of human lungs. RNA-seq data from the Genotype-Tissue Expression (GTEx) project was analyzed to determine gene expression changes in non-diseased lung tissue from various ages. Dataset from young (subgroups 20-29, 30-39, 40-49) was compared with elders (subgroups 50-59, 60-69, 70-79). A) Gene Set Enrichment analysis (GSEA) of inflammatory pathways enriched in elderly relative to young. B) GSEA of ECM related/remodeling pathways over-represented in elderly compared to young data sets. C) xCell analysis comparing young and elders showing the most abundantly expressed cell types. D) xCell analysis comparing young and elders showing cell types of medium expression. E) xCell analysis comparing young and elders showing levels of cell types with the lowest expression. Pairwise comparison with unpaired two-samples Wilcoxon test (*P ≤ 0.05, **P ≤ 0.01, ***P ≤ 0.001).

With the young and old lung transcriptomes, we used a deconvolution method xCell^29^ to enumerate different cell types based on the gene expression profiles. Of the most abundant cell types represented, we identified that CD4 memory T cells, monocytes, and mesenchymal stem cells were significantly increased in the older compared to the young lungs, while total macrophages and activated dendritic cells (aDCs) were decreased in the old lungs (Fig 1C). Of the medium abundant cell types, CD4^+^ T cells, CD8^+^ T cells and regulatory T cells (Tregs), as well as B cells and neutrophils were over-represented in the older lungs relative to the young (Fig 1D). Of the low abundant cell types, lymphocytes, natural killer (NK) cells, gamma-delta T cells, and platelets were also significantly up in the older lungs (Fig 1E). Based on these analyses of the transcriptomes from human lung tissue, there is considerably alterations in the tissue microenvironment marked by remodeling of the ECM and differences in the immune landscape with old age.

### Aging contributes to cellular and phenotypic changes in mouse lungs

We next asked whether similar age-dependent transcriptional changes occur in the lungs of young compared to aged mice. We mined gene expression datasets analyzed via Affymetrix 5 of purified lung epithelial tissue isolated from 3-month and 18-month-old mice^28^. Pathway analyses revealed numerous pathways under-represented (Fig. S2A) and over-represented (Fig. S2B) in old lung epithelium compared to young. Of the top 20 most significant pathways, alterations in the extracellular matrix and tissue structure such as ECM-receptor interaction, tight junction, regulation of cytoskeleton, tissue morphogenesis, and actin filament-based process were significantly down-regulated in old lung epithelium compared to young (Fig. 2A). These results suggest there are structural changes and remodeling of the lung tissue epithelium which occur with old age. Conversely, when we analyzed pathways upregulated in aged epithelium, we found an upregulation of cilium related pathways including cilium assembly, cilium dependent cell mobility and protein localization to cilium included in the top 20 most significant pathways (Fig. 2B). Previous single-cell RNA-seq studies have demonstrated an increase in the frequency of ciliated cells in the lungs of aged mice compared to young^37^. Additionally, we observed an increase in immune and inflammation related pathways such as hematopoietic cell lineage and chemokine signaling pathway in lung epithelium of aged mice (Fig. 2B).

**Figure 2.**
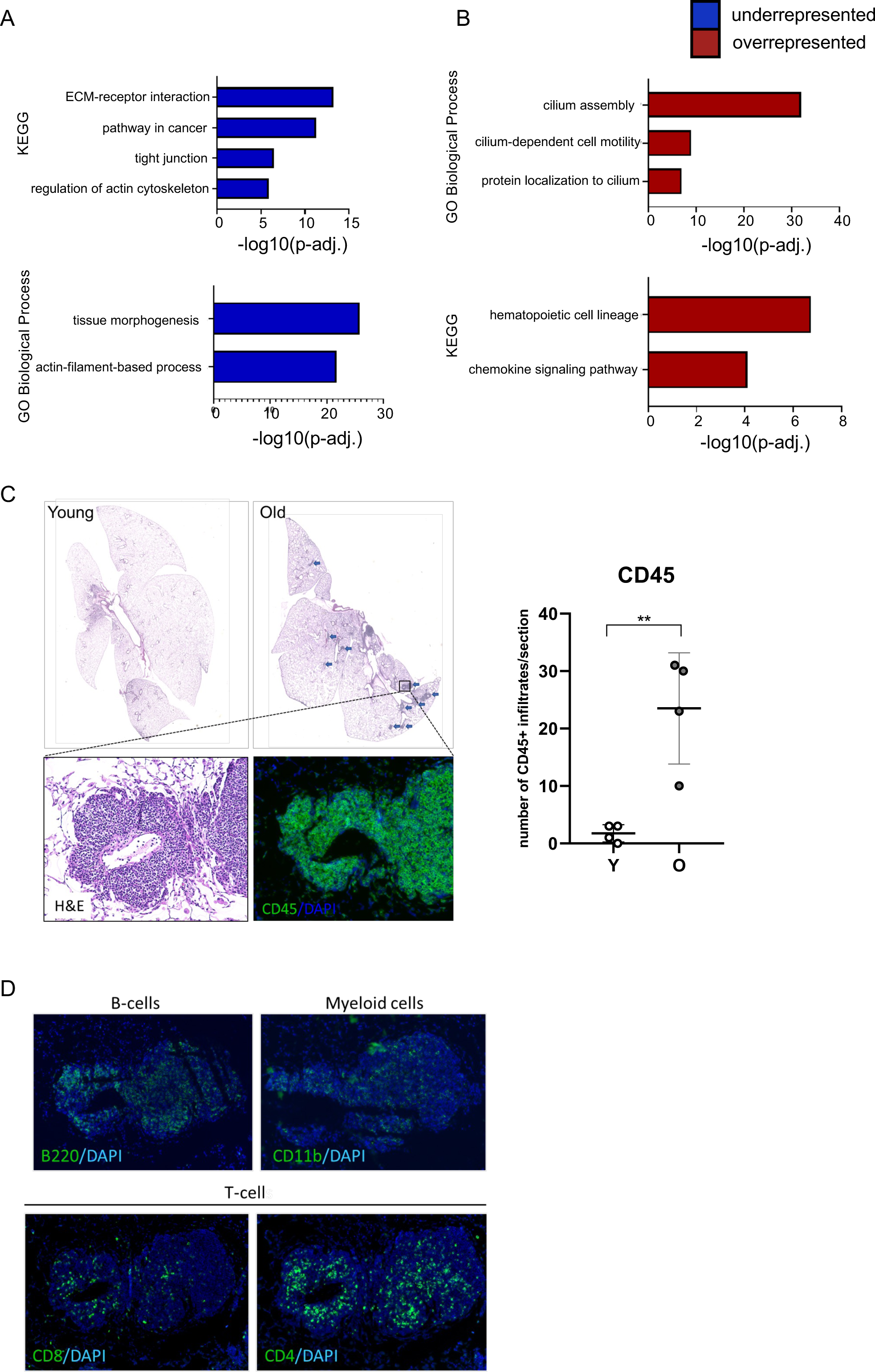
Aging contributes to cellular and phenotypic changes in mouse lungs. Gene expression datasets analyzed via Affymetrix of lung epithelial tissue isolated from 3-month and 18-month-old mice were mined. Metascape analysis was performed on the most significantly differentiated genes in epithelial cells of lungs from young and old mice. A) Pathways associated with ECM and remodeling under-represented in old epithelium compared to young. B) Pathways over-represented in old epithelium compared to young. C) Lungs from young (2-month-old) and old (20-month-old) mice were harvested and inflated for histology. Leukocytic marker CD45 was stained using immunofluorescence to quantify immune lesions in the lungs of young and old mice. D) Additional immunofluorescence analysis was performed to probe for B220 (b-cell marker), CD11b (myeloid marker), CD4 (helper T-cell marker) and CD8 (cytotoxic T-cell marker).

To determine whether there were cellular and structural changes associated with old age in the lungs, we harvested lungs from young mice (2 months) and old mice (20 months). Unlike lungs from young mice, lungs from old mice appeared to have an increase in the number of immune aggregates as indicated by staining of pan-leukocyte marker CD45 compared to the lungs of young counterparts (Fig. 2C). These immune aggregates were observed in close proximity to the bronchial space as well as throughout the lung parenchyma. In order to identify the type of immune cells that make up the immune lesions present in old lungs, we performed immunofluorescence and stained for cd11b (myeloid marker), B220 (B-cell marker) and CD4/8 (T cell markers).

We found significant positivity of B220 expression and to a lesser extent, CD11b (myeloid marker) positivity within the immune aggregates (Fig. 2D). Furthermore, we identified the presence of both CD4 and CD8 T cells, also within the immune aggregates (Fig. 2D). It has been reported that the adaptive immune system exhibits several age-associated changes including an increase in the number of effector and memory CD4^+^ T-cells^38^. These immune aggregates identified in aged lungs resemble immune lesions such inducible bronchus-associated lymphoid structures (iBALTs) found in smoking lungs^39^. Based on the mouse and human data, our results suggest that there is an age-associated increase in immune infiltrates in the lung.

### Expression of AAT reduces inflammatory phenotype and partially reverse age-dependent gene expression alterations in aged lung

Given that we observed an increase in immune infiltrates and an enrichment in inflammatory pathways in our gene expression analysis in the lungs of aged mice, we sought out a model that could possibly dampen these effects of aging in the lung. We have previously shown that aged AAT transgenic mice reversed increased TNF-α in serum and bone marrow and IL-6 in bone marrow of old mice^40^. Additionally, AAT mice have been reported to be resistant to several inflammatory challenges^41^. The human AAT transgene is expressed from the surfactant protein C (SPC) promoter^23^; while a secreted protein, we expect higher levels in the lungs. Interestingly, we detected far fewer immune aggregates in old-AAT mice, similarly to the lungs of young mice (Fig. 3A). To gain mechanistic insight for how AAT expression may be impacting age-dependent alterations in the lungs, we performed RNA sequencing analysis of whole lung to evaluate gene expression profiles among the different groups. We observed a significant proportion of genes altered in young compared to old lungs (Supplemental Table 1, 2). Interestingly, AAT expression in old mice was able to partially reverse some of these changes. We identified 153 genes up in young mice relative to old and 342 genes up in old relative to young. We also saw 53 genes up in old relative to old-AAT lungs and 298 genes up in old-AAT in comparison to old lungs.

**Figure 3.**
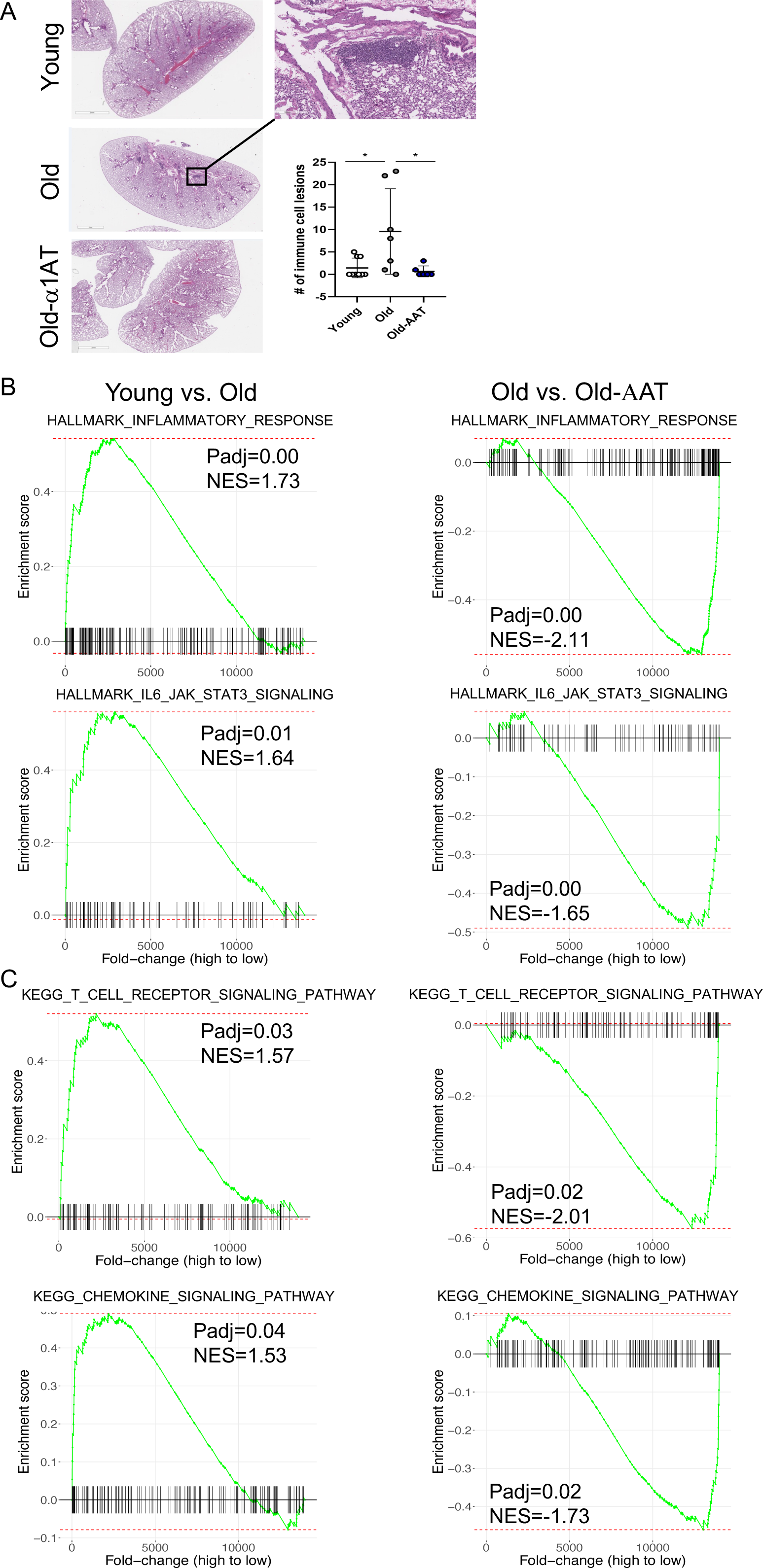
Expression of AAT reduces inflammatory phenotype and partially reverse age-dependent gene expression alterations in aged lung. A) The number of distinct immune areas (as shown) were quantified in young, old and old-AAT mice (ANOVA; *P ≤ 0.05, **P ≤ 0.01, ***P ≤ 0.001). Functional analysis of RNA-sequencing of lungs from young, old, and old-AAT mice. Pathways identified as being over-represented in old and restored to youthful levels with AAT expression. B) Hallmark pathway analysis showed upregulated inflammatory response and IL6/JAK-STAT3 signaling in old mouse lungs, and a reversal with AAT overexpression. C) KEGG pathways analysis showed upregulated T cell receptor signaling pathway and chemokine signaling pathway in old mouse lungs, and a reversal with AAT overexpression.

We next performed gene set enrichment analysis (GSEA) on the significant genes identified in young, old, and old-AAT groups. From the Hallmark database, we identified inflammatory response and IL-6/JAK/STAT3 signaling enriched in old relative to young, with the opposite enrichment with AAT expression, indicating reversal of this aging-dependent pathway upregulation (Fig. 3B). We also identified T cell receptor signaling pathway and chemokine signaling pathway to be up in old lungs and down with AAT expression from the KEGG database (Fig. 3C). As for the IL-6/JAK/STAT3 pathway, expression of AAT restores expression of these T cell and chemokine pathway genes to more youthful levels. Lastly, we identified PD-1 signaling pathway being more activated, with AAT expression reversing the activation (Fig. S3A). Flow cytometric analyses of CD4 and CD8 T cells similarly revealed an increased PD1^+^CD4^+^ and PD1^+^CD8^+^ T cells in old lungs compared to young ones, with a reversal with AAT overexpression (Fig. S3B). Nonetheless, we did not observe reductions in the aging-associated increased adenoma outgrowth upon injection of anti-PD1 (Fig S3C), previously shown to promote immune reactivation^42^.

### Expression of AAT partially restores tissue microenvironmental alterations in aged lung

We next employed a proteomics approach to ask how aging may impact the architecture of the tissue microenvironment and how AAT expression may affect these changes in the lungs. To address this, we performed mass spectrometry (MS)-based proteomic analyses of lung tissue from young, old, and old-AAT mice. We were able to evaluate changes from the whole proteome, as well as the extracellular matrix (ECM) that contributes substantially to the tissue architecture of the microenvironment^27^. Lung samples from young, old, and old-AAT mice were separated into high salt (HS, enriched for cellular proteins), soluble ECM (sECM), and insoluble fractions (iECM) (Fig 4A). We assessed global proteomic changes and found dramatic alterations among young and old lungs. Interestingly, AAT was able to mitigate a subset of those changes among the different fractions (Fig. 4B). Both of the tight junction proteins ZO-1 and ZO-3 were down in old lungs and effectively restored in old lungs expressing AAT in the iECM and sECM, respectively. Furthermore, the distribution of ZO-1 was changed substantially, with reduced presence in the iECM and increases in HS fraction in the lungs of old mice, suggesting age-associated tissue remodeling. In addition, these analyses revealed significant increases in the levels of neutrophilic proteases (e.g. MMP9 and ELNE/elastase) and other inflammatory markers (e.g. S100A8/A9) in the old lungs, as well as changes in key factors dictating cell fate (e.g. NEST/Nestin), which were effectively reversed by AAT overexpression (Fig. 4C). Pathway analysis of all significant protein changes further showed an increase in ECM remodeling (ie. cell-cell adhesion and actin cytoskeleton remodeling) and inflammation (i.e. Inflammatory response, leukocyte activation, IL-17 signaling) enriched in old lungs and restored to youthful levels with AAT expression (Fig. 4D). Additionally, indications of ECM remodeling were also evident by over-representation of protein folding and actomyosin structure organization, and tight junction pathways that were under-represented in old lungs and enriched in old-AAT lungs (Fig. 4E). In all, the proteomic analyses reveal substantial changes in the tissue microenvironment with age, which can partially be restored with AAT expression.

**Figure 4.**
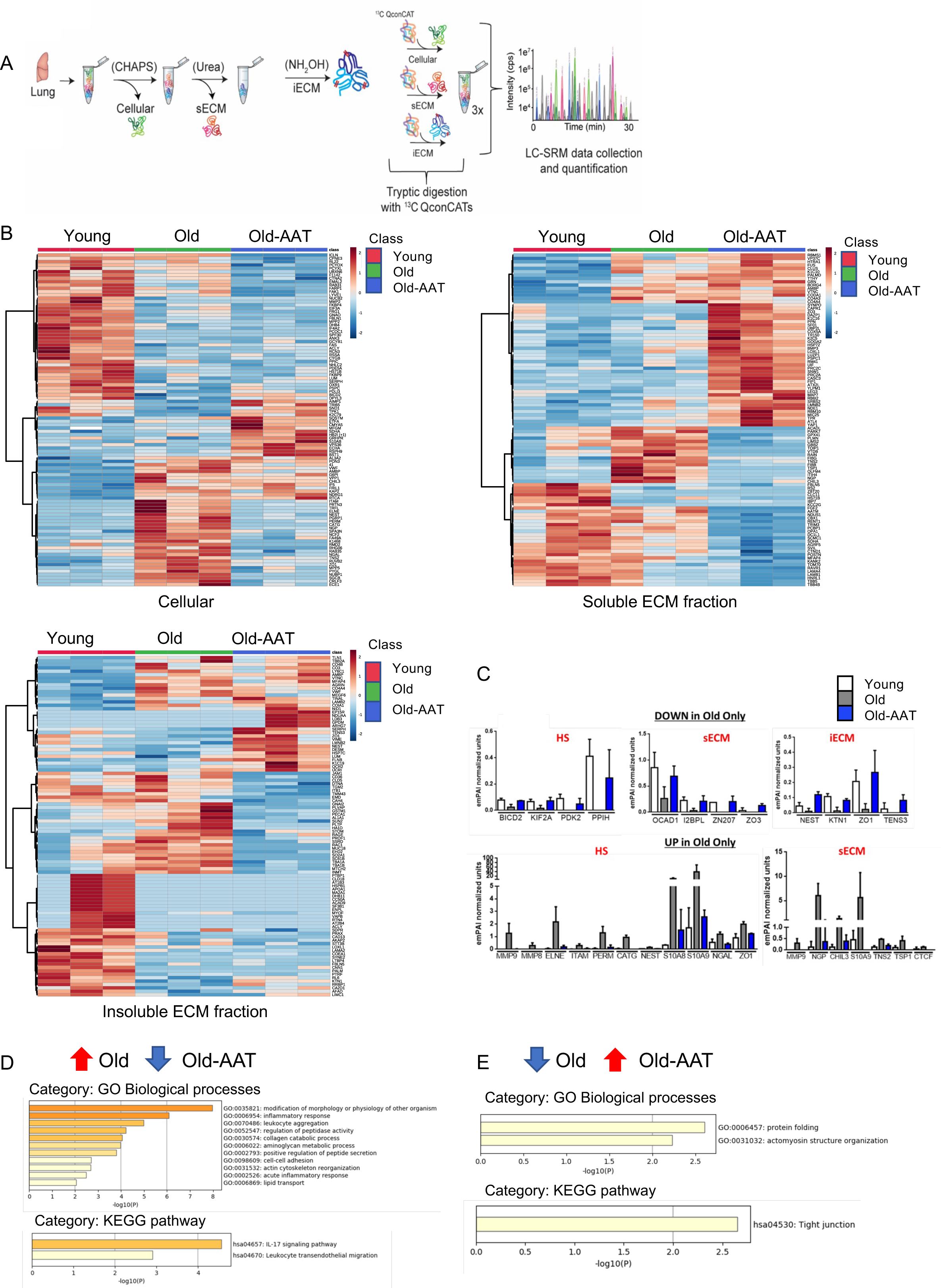
Expression of AAT partially restores tissue microenvironment alterations in aged lung. A) Schematic of extraction of high salt (HS), soluble ECM (sECM) and insoluble ECM (iECM) fractions for mass spectrometry analysis. Lung tissues from young (3 months), old (24 months) and old-AAT (24 months) mice were analyzed. B) Heat map of HS, sECM, and iECM of proteomic changes with young, old and old-AAT lung tissue. C) The top row selected proteins reduced in aged lungs but not in young or old-AAT. The bottom row selected proteins increased in old lungs. All graphed changes were significant by ANOVA (p<0.05). EmPAI units normalized the number of detected peptides to the total number of detectable peptides per protein, and then to total protein content per sample. Metascape analysis was performed using D) lists of significantly up-regulated proteins in old lungs which were significantly down-regulated with AAT expression E) lists of significantly down-regulated proteins in old lungs which were significantly up-regulated with AAT expression.

### Age-dependent increased inflammatory parameters are also observed in lung adenocarcinomas and squamous cell carcinomas

Given the changes observed in aging lungs, we wondered if similar changes occur in lung cancer tissue dependent on patient age. We thus analyzed the publicly available RNA-seq data from The Cancer Genome Atlas (TCGA) of over 400 lung adenocarcinomas or lung squamous cell carcinomas. We similarly performed GSEA analyses comparing gene expression profiles of lung cancer tissues from young (40-49) and old (70-79) patients with lung adenocarcinoma and from young (50-59) and old (70-79) patients with lung squamous cell carcinoma. Similar to normal aging lungs (Fig. 1A, S1A), pathways related to inflammation, including inflammatory response, TNFA signaling via NFκB, interferon gamma response were shown to be upregulated in elderly patients with both lung adenocarcinoma and lung squamous cell carcinoma (Fig. 5A, B), suggesting that old age confers its inflammatory nature to cancers that develop within the lungs. Myc targets and oxidative phosphorylation gene sets were also identified to be downregulated in both cancer types with age (Fig. 5C, D), similar to our observations in normal aging lungs (Fig. S1B), suggestive of a common force at play in lung cancer development.

**Figure 5.**
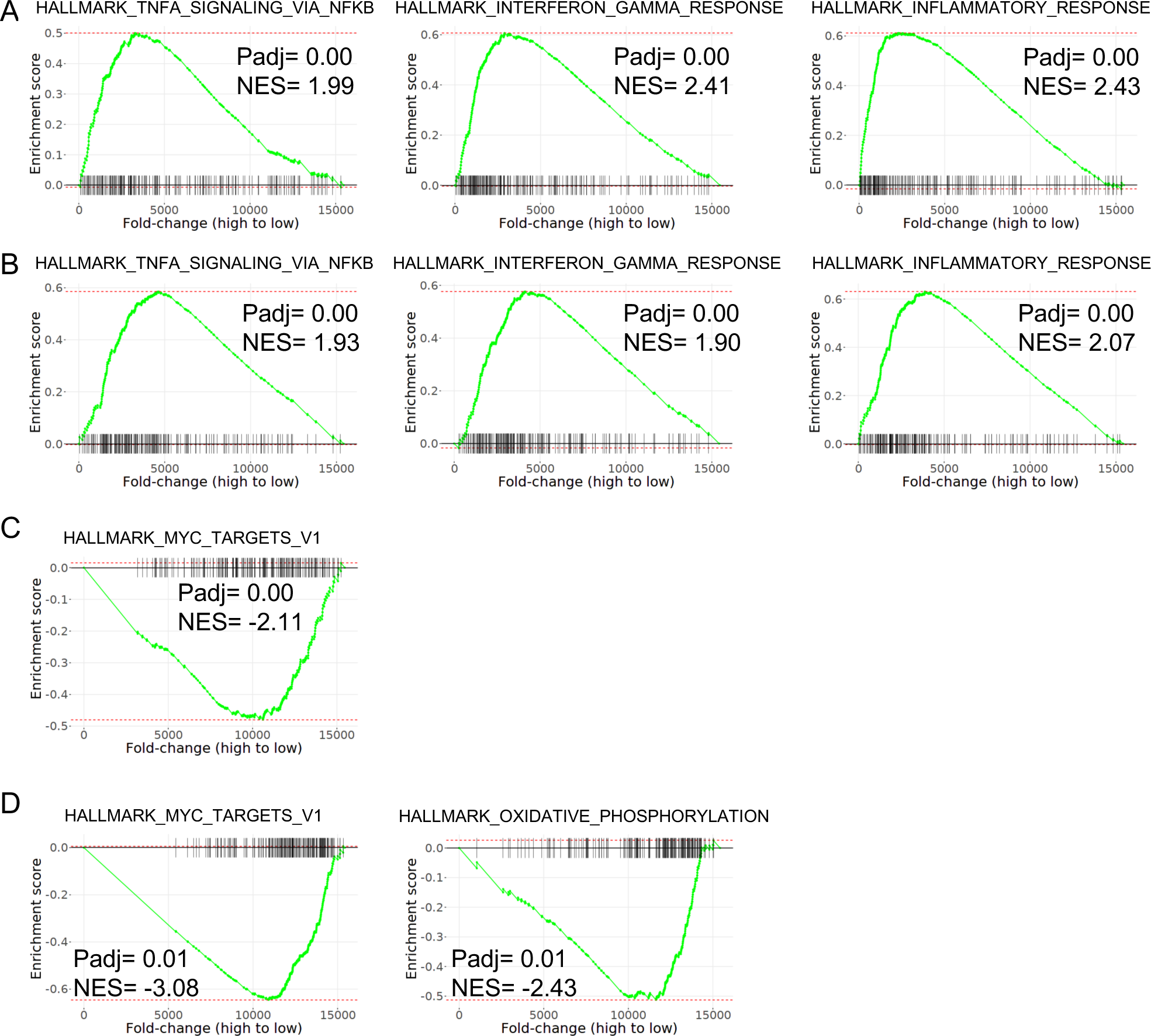
GSEA analyses of the Genome Atlas (TCGA) RNA-seq data from lung adenocarcinoma and squamous cell carcinoma patients revealed a more inflamed microenvironment in elderly patients. A) Hallmark pathway analyses showed upregulated TNFA_Signaling_Via_NFκB pathway, Interferon_Gamma_Response, and Inflammatory_Response in lung adenocarcinoma patients aged 70-79 compared to those of 40-49. B) Hallmark pathway analyses showed upregulated TNFA_Signaling_Via_NFκB pathway, Interferon_Gamma_Response, and Inflammatory_Response in lung squamous cell carcinoma patients aged 70-79 compared to those of 50-59. C, D) Hallmark pathway analyses showed downregulated MYC_TARGETS_V1 in lung adenocarcinoma patients, MYC_TARGETS_V1 and OXIDATIVE_PHOSPHORYLATION in lung squamous cell carcinoma patients with the same age-group comparison in A and B.

### Age-dependent selection of oncogenic initiated clonal expansions in lung mouse model

To determine whether aging-associated alterations in the tissue microenvironment could be conducive to oncogenic adaptation, we used an orthotopic model whereby cells derived from EML4-ALK fusion-induced adenomas (EA cells) are directly implanted into lungs of immunocompetent young and old mice^25^. Note that the EA cells were generated in young mice, and thus we are only assessing the impact of the lung environment of the recipient mice. Using this model, we demonstrated an increased tumor growth in lungs of old mice compared to young mice (Fig. 6A, B). NLR family pyrin domain containing 3 (NLRP3) inflammasome is a protein complex that mediates the activation of inflammatory mediators^43^. NLRP3 is required for the production of mature IL-1β and IL-18^43^. Given that transgenic AAT expression reversed some of the aging-dependent changes in lungs, including increased inflammatory response (Fig. 1A, S1A) and that both AAT deficiency and NLRP3 inflammasome contribute to lung inflammation through similar pathways^44,45^, we asked whether NLRP3 deficiency could prevent increased EA cells outgrowth in old mice. We implanted EA cells directly into lungs of young, old, and old *Nlrp3^-/-^* mice, and observed a subdued EA cells outgrowth in old *Nlrp3^-/-^* mice compared to wild type old mice (Fig. 6B). While we no longer observed increased tumorigenesis in the aged *Nlrp3^-/-^* mice compared to young mice, we note that the difference between old wild-type and old *Nlrp3^-/-^* mice was not significant. Thus, the aged lung microenvironment promotes the expansion of adenoma cells, and this effect appears dampened by the loss of IL-1β production.

**Figure 6.**
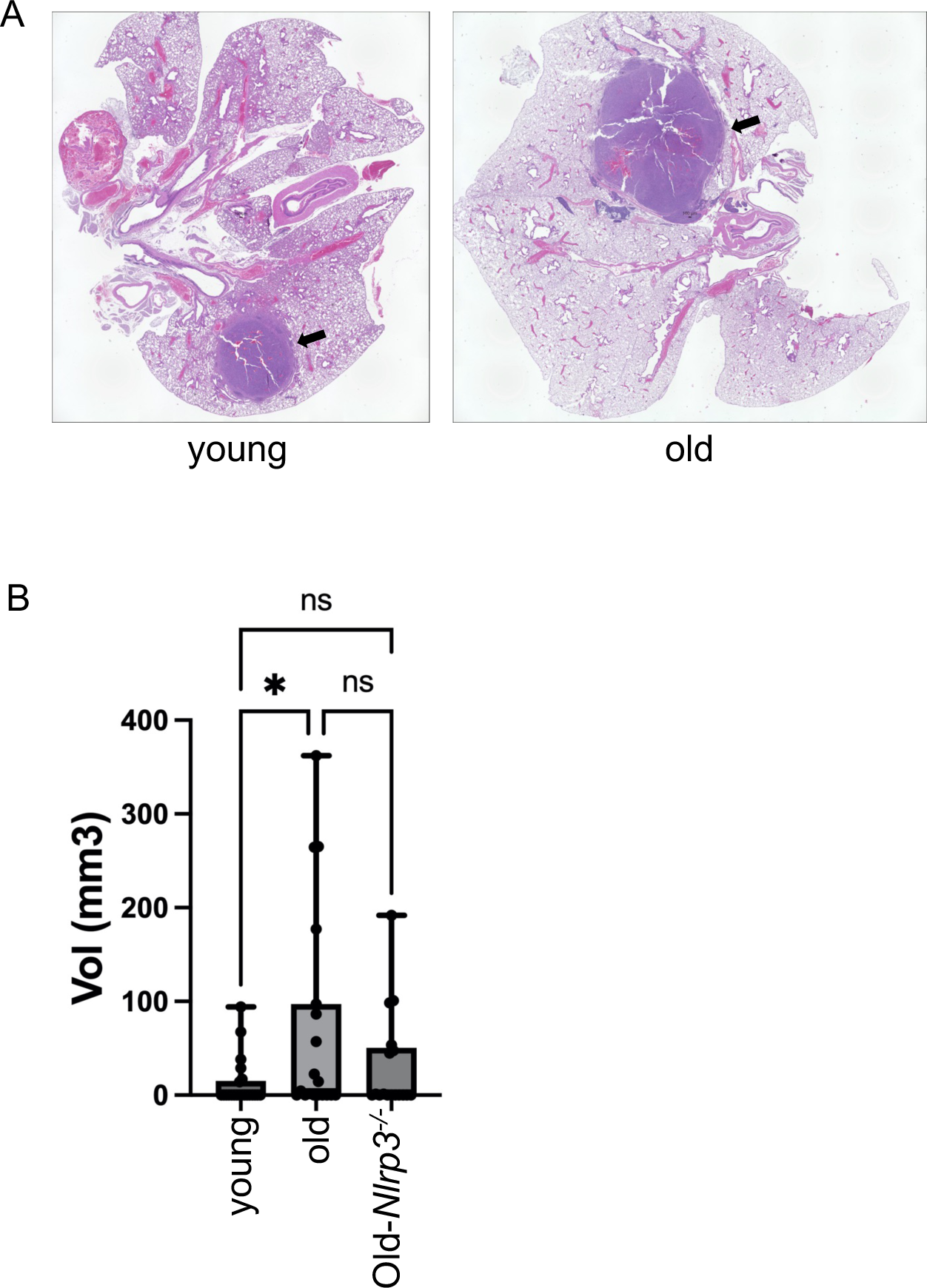
Age-dependent selection of oncogenic clonal expansions in lung mouse model. 2×10^5^ EML4-ALK cells per mouse were implanted into the left lungs. Orthotopic tumors established for 4 weeks where mice were imaged using μCT imaging to determine tumor volumes before harvesting. A) Images of orthotopic tumors from young and old mouse lungs 4 weeks after the initial implantation. B) Tumor volumes (mm^3^) comparing tumor growth from lungs of young, old, and old-*Nlrp3^-/-^* mice.

## DISCUSSION

While it is widely recognized that mutations are crucial for the development of cancer, the role of the microenvironment and importantly the notion that selection of particular mutations is context-dependent are often underappreciated in the field of tumorigenesis. Previous work has demonstrated that under the contexts of microenvironmental insults such as radiation and aging can promote oncogenic selection in the hematopoietic system^18,46,47^. The results presented here further demonstrate that aging-associated inflammation can impact tumor development driven by oncogenic EML4-ALK in the lungs.

Lung cancer, like most other cancers, is an aging-related disease. Aging results in an inflammaging phenotype in the lungs marked by increases of immune cell infiltrates, predominately but not limited to cells involved in the innate immune system^48^. It has previously been shown that in a conditional mouse model of oncogenic Kras^G12D^, activation of Kras^G12D^ in the lungs led to higher numbers of adenomas in old mice compared to their young counterparts^49^. Based on gene expression analyses of initiated tumors in young and old mice, the authors speculated that tumor formation in the aged lung was in part attributed to attenuated DNA damage and p53 tumor repressor responses, as well as an increased inflammatory milieu. Our data provide evidence for a causative link between the aged microenvironment, increased inflammation, and cancer susceptibility.

Our studies characterized how aging changes the immune landscape of lungs. We also observed changes in the immune landscape of aged lungs compared to young human lungs through mining the GTEx database. Inflammatory pathways including pathways related to T cell activation and signaling are enriched in aged lungs relative to young. Using xCell, we observed significant increases in the number of CD4^+^ and CD8^+^ T cell subtypes, notably an increase in memory T cells. Moreover, aged lungs displayed increases in Tregs and neutrophils, alterations in macrophage subsets, and decreases in dendritic cells. The dysregulation of both innate and adaptive immunity has shown in the lungs of aged individuals^50^. In the elderly, pro-inflammatory cytokines have been shown to be significantly elevated with an increase in immune cell infiltrates involved in innate immunity^51^. Specifically, increases in IL-1β, IL-6 and TNF-α have been observed^52,53^. This phenomenon occurs in the apparent absence of an immunologic threat or infection, which may contribute to the serum proteinase-mediated weakening of the lung parenchyma with old age. In addition, adaptive immunity has also been shown to decline with old age including alterations in the number of naïve and memory CD4^+^ and CD8^+^ T-cells as well as B cells^54,55^. Systemic increases of Tregs have also been reported with aging^56^, further contributing to enhanced susceptibility to infection in the elderly.

In comparing mouse lung tissue to human, we observed similar age-dependent changes in hallmarks of inflammation. Through analyses of microarray data from young and old mouse epithelium, inflammatory (hematopoietic cell lineage, chemokine signaling pathway) and ECM remodeling-related pathways (tight junction, regulation of actin cytoskeleton) were significantly changed in young versus old epithelium, suggesting that the epithelial cells have intrinsic alterations, like the bulk lung tissue.

Notably, when we perform immunofluorescence, we observe an increase in immune aggregates in aged lungs, with a major component of T cells. These immune lesions are reminiscent of tertiary lymphoid structures, which have been reported to positively correlate with immunotherapy in cancers^57,58^; however, the role of these immune aggregates with old age and cancer initiation has yet to be elucidated. Due to the important role of inflammation in cancers^59^ and from the inflammatory phenotype in aged lungs observed from our studies, we asked whether aging-associated inflammation may be contributing to oncogenic selection. To this end, we utilized an anti-inflammatory mouse model with transgenic expression of human AAT. Expression of AAT in aged lungs significantly reduced the number of immune aggregates relative to aged, similar to that of young lungs.

Through RNA sequencing, we identified pathways related to inflammation that were upregulated in old lungs, but restored or partially restored to more youthful levels in old lungs expressing AAT. These pathways included inflammatory response, innate immune response, cytokine activity and chemokine signaling pathways. Upon closer inspection, we found that chemokines including CCL8, and CCL5 were up in old and restored to levels observed in young in the lungs of old-AAT mice. Interestingly, CCL5 has been shown to play a role in the recruitment of neutrophils specifically in the lungs^60^. Additionally, CCL5 and CCL8 have been reported to be chemoattractants for many immune cell types including monocytes, eosinophils and lymphocytes^61^. Future studies will need to explore the possible roles for these and other chemokines in promoting the aging-associated immune cell influx in the lungs of older individuals (whether mouse or human). The ability of AAT expression to prevent some of these immune changes opens up the possibility of developing interventions to mitigate some of these aging-associated changes, with the potential to lessen lung cancer risk.

We also performed proteomics to assess alterations in the ECM of the lung tissue microenvironment. We observed dynamic changes in the ECM as seen through the redistribution of tight junction protein, ZO-1, in old lungs which was prevented with expression of AAT. The ECM not only serves as a scaffold for the overall tissue architecture, but also serves as a niche that is crucial for determining cell fate^62^. Additionally, key proteases such as MMP9 and ELNE were also found to be elevated in the HS fraction containing cellular proteins specifically in old lungs and restored to youthful levels with AAT expression. A previous study showed notable changes in inflammatory genes (*Il6*, *Il1b*, *Tnf*, *Il21*, *Il5*) at single cell resolution in the mouse lung^37^. In accordance with these observations, we observed significant differences in inflammatory markers (e.g. S100A8/A9) in old lungs compared to both young and old-AAT indicating that inflammation is a prominent phenotype in aged lungs. Despite being able to separate the HS fraction containing cellular proteins and both the insoluble and soluble ECM fractions, the ECM is complex in its composition and posttranslational modifications. Future studies will be required to determine whether there are differences in protein crosslinking with age, and whether expression of AAT impacts this crosslinking. It is notable that only a subset of the mRNA and protein changes observed in old lungs were reversed by AAT expression. Taken together, our data demonstrate that the there is considerable age-dependent remodeling of the tissue architecture of the lung. Furthermore, expression of AAT was able to prevent some of these age-dependent changes.

Future studies will be required to investigate the role of key mediators during inflammation, such as IL-1α, TNF-α and IL-6, and whether inhibition of these cytokines in aged mice is sufficient to limit oncogenesis in the lungs. The availability of senolytic therapies (ie. ABT-263)^63^ or anti-inflammatory agents (NLRP3 inhibitor, OLT1177)^64^ could also potentially serve as therapeutics to modulate the tissue microenvironment to dampen oncogenic adaptation. Since the process of aging is complex, it is unlikely that changing one aspect of aging could completely prevent oncogenesis; however, it is possible to identify significant contributors involved in oncogenic adaptation. In all, our studies have revealed marked changes in the lung ECM remodeling and inflammation that accompany aging, and demonstrate how these changes persist in lung cancers and likely contribute to lung cancer evolution. Importantly, while we are unlikely to be able to prevent many of the mutations that accumulate in our tissues during aging, thus contributing to cancer evolution, we do have the means to manipulate tissue microenvironments to alter cancer evolutionary trajectories.

## Acknowledgements

These studies were supported by grants from the Veteran’s Administration to J.D. (1 I01 BX004495). H.A.S. was supported by T32CA174648. E.A.B. was supported by University of Vermont College of Medicine New Research Initiative Grant. The authors thank the Bioinformatics and Biostatistics, Genomics, Mass Spectrometry and Flow Cytometry Shared Resources supported by National Cancer Institute grant P30CA046934 to the University of Colorado Cancer Center.

---

**Supplemental figure 1.**
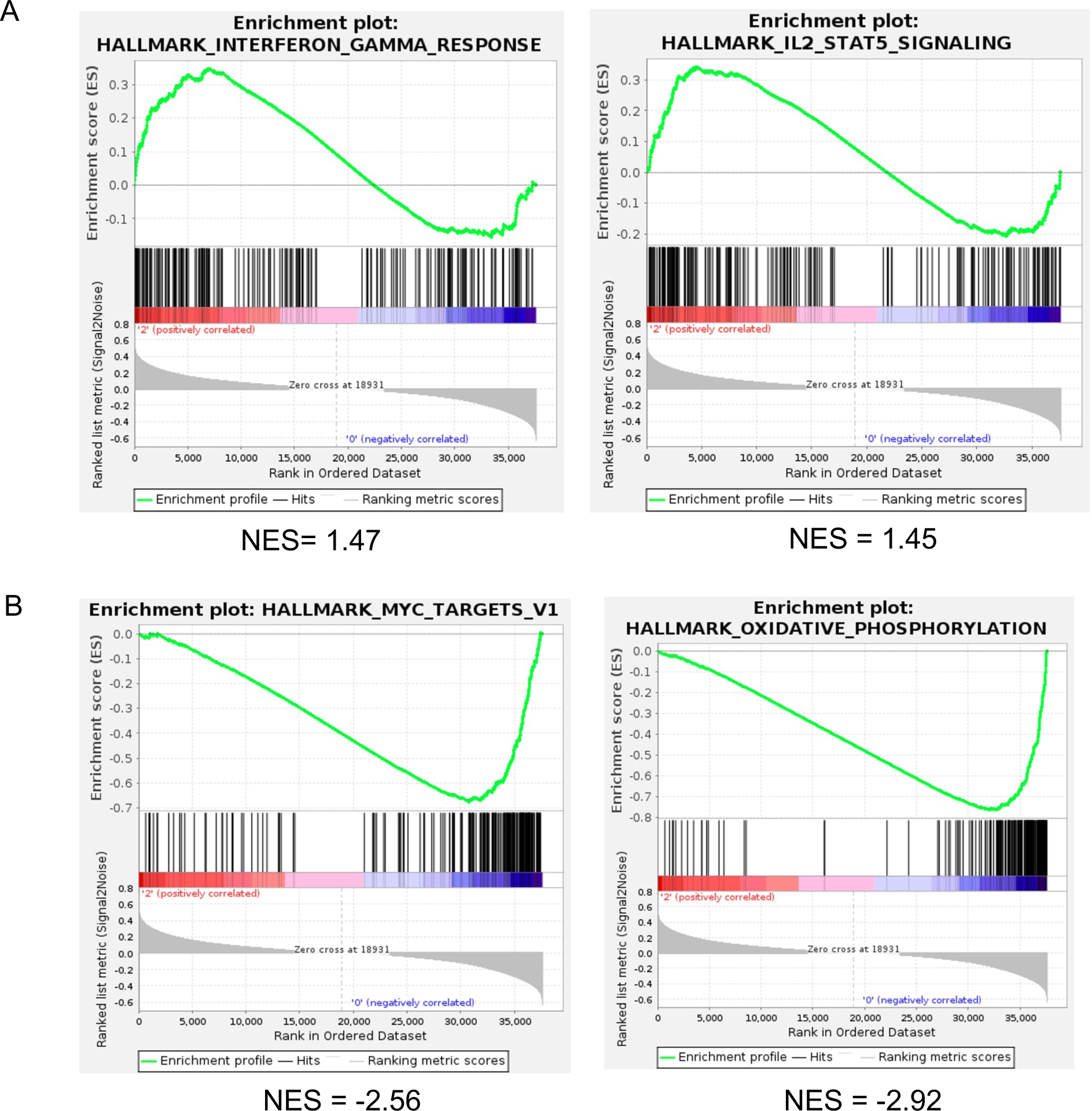
GSEA analysis reveals age associated alterations in pathways and cell types. A) Interferon gamma response and IL2-STAT5 signaling pathways are enriched in the lungs of elderly versus young non-diseased lungs using GSEA of GTEx project. B) Hallmark Myc targets and oxidative pathways under-represented in elder in comparison to young lungs.

**Supplemental Figure 2.**
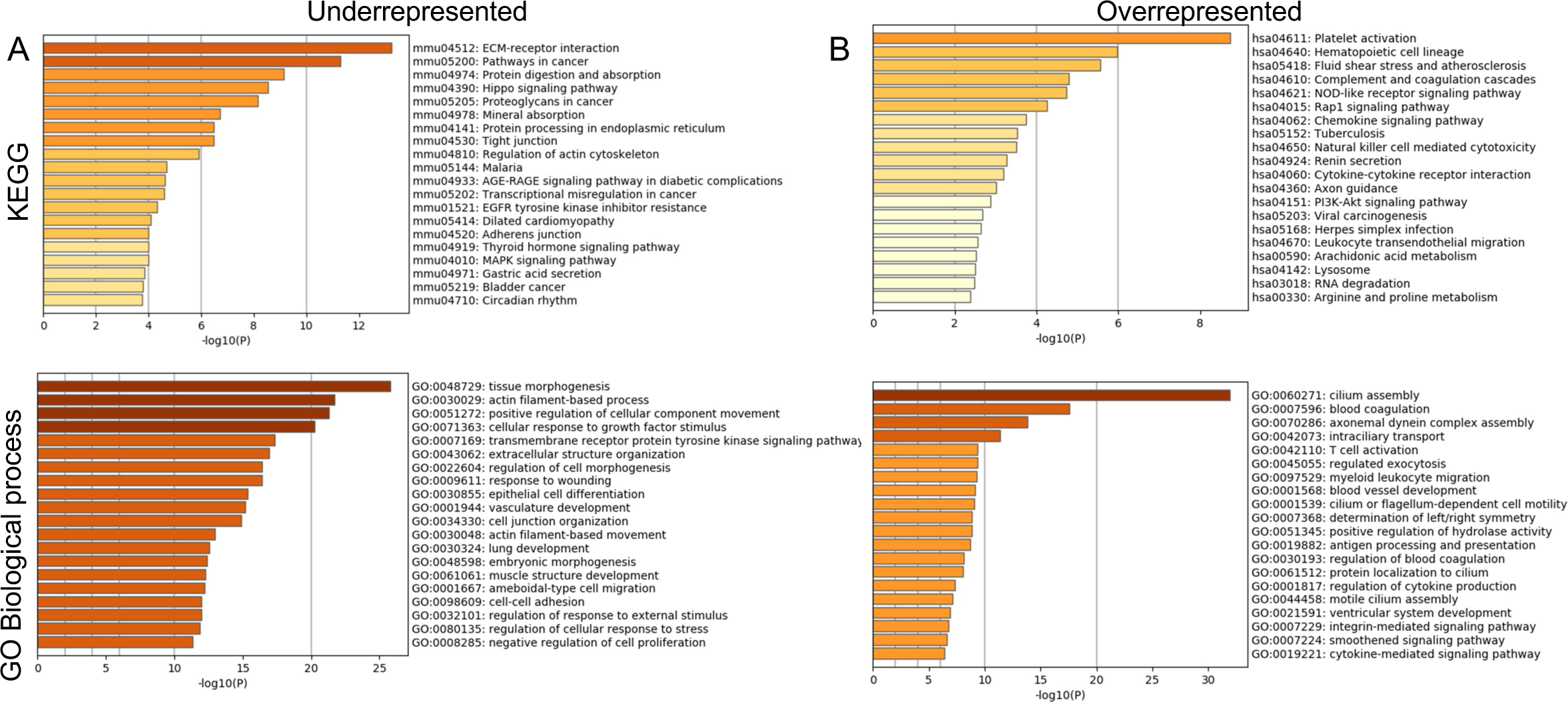
Age-dependent changes are observed in lung epithelium. Metascape analysis was performed on the most significant genes found in young versus old lung epithelium tissue. A) Representation of the top 20 most significantly under-represented in old epithelium compared to young from KEGG and GO biological process datasets. B) Representation of the top 20 most significantly over-represented in old epithelium compared to young from KEGG and GO biological process datasets.

**Supplemental Figure 3.**
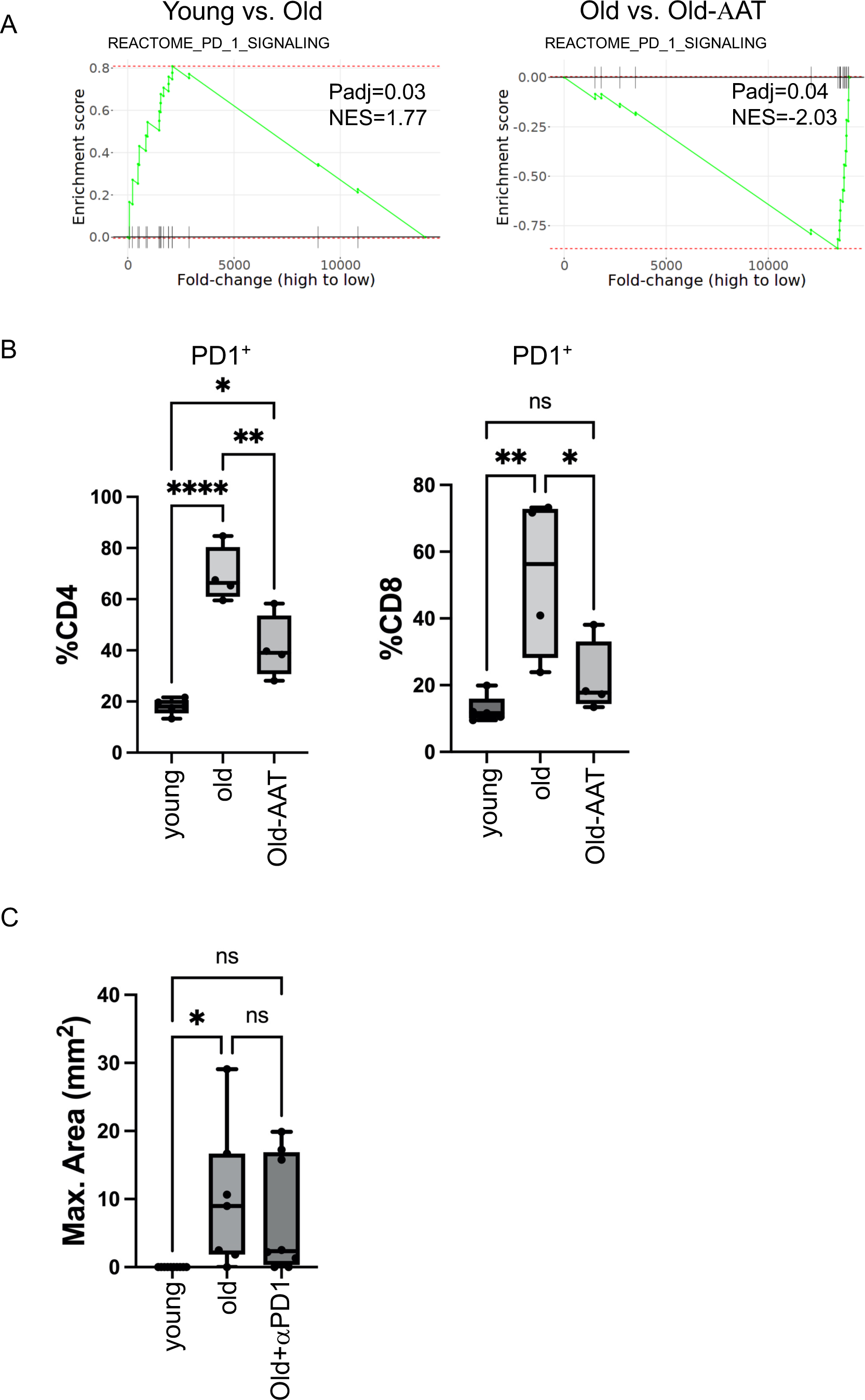
Expression of AAT reversed age-dependent increased activation of PD-1 signaling pathway. A) Reactome pathway analysis showed upregulated PD-1 signaling in old mouse lungs, and a reversal with AAT overexpression. B) Flow cytometric analyses of CD4 and 8 T cells showing increased percentage of PD1^+^ CD4 and 8 T cells, and a reversal with AAT overexpression. C) PD1 blocking antibody was injected 2× per week, starting 2 weeks before 2×10^5^ EML4-ALK cells per mouse were implanted into the left lungs. Lungs were harvested 4 weeks after the implantation and stained with hematoxylin and eosin. Maximum areas were identified through serial sections of the entire lungs and measured using ImageJ. Tumors displayed larger areas in aged mice compared to tumors in young mice, while PD-1 blocking does not reverse the increased tumor growth in aged mice.

